# FLN-2 functions in parallel to LINC complexes and Cdc42/actin pathways during P-cell nuclear migration through constricted spaces in *Caenorhabditis elegans*

**DOI:** 10.1101/2023.08.04.552041

**Authors:** Linda Ma, Jonathan Kuhn, Yu-Tai Chang, Daniel Elnatan, G.W. Gant Luxton, Daniel A. Starr

## Abstract

Nuclear migration through narrow constrictions is important for development, metastasis, and pro-inflammatory responses. Studies performed in tissue culture cells have implicated LINC (linker of nucleoskeleton and cytoskeleton) complexes, microtubule motors, the actin cytoskeleton, and nuclear envelope repair machinery as important mediators of nuclear movements through constricted spaces. However, little is understood about how these mechanisms operate to move nuclei *in vivo*. In *C. elegans* larvae, 6 pairs of hypodermal P cells migrate from lateral to ventral positions through a constricted space between the body wall muscles and the cuticle. P-cell nuclear migration is mediated in part by LINC complexes using a microtubule-based pathway and by an independent CDC-42/actin-based pathway. However, when both LINC complex and actin-based pathways are knocked out, many nuclei still migrate, suggesting the existence of additional pathways. Here we show that FLN-2 functions in a third pathway to mediate P-cell nuclear migration. The predicted N-terminal actin binding domain in FLN-2 that is found in canonical filamins is dispensable for FLN-2 function, this and structural predictions suggest that FLN-2 is not a divergent filamin. The immunoglobulin (Ig)-like repeats 4-8 of FLN-2 were necessary for P-cell nuclear migration. Furthermore, in the absence of the LINC complex component *unc-84*, *fln-2* mutants had an increase in P-cell nuclear rupture. We conclude that FLN-2 functions to maintain the integrity of the nuclear envelope in parallel with the LINC complex and CDC-42/actin-based pathways to move P-cell nuclei through constricted spaces.

## Introduction

The positioning of nuclei is important to many developmental processes, including pronuclear migration, muscle development, and neurodevelopment (Bone and Starr 2016). However, moving nuclei to the proper subcellular location, especially if it needs to traverse a constricted space, is poorly understood. Nuclei are the largest and stiffest organelle, making nuclear deformation and migration the rate limiting step for cell migrations through constricted spaces significantly smaller than the resting diameter of nuclei (Davidson *et al*. 2014). For example, during hematopoiesis, white blood cell nuclei migrate out of the bone marrow, while the stiffer nuclei of megakaryocytes cannot, leading to the formation of anucleated platelets (Junt *et al*. 2007; Shin *et al*. 2013; McGregor *et al*. 2016; Salvermoser *et al*. 2018). Metastasis of some cancer cells also relies on the deformation of their nuclei so that the cells can squeeze through small opening in the extracellular matrix (Seyfried and Huysentruyt 2012; Fu *et al*. 2012; Gensbittel *et al*. 2021). Thus, understanding how nuclei normally traverse constricted spaces could help our understanding of metastasis. Tissue cultured cells can be studied migrating through fabricated constrictions as small as 5% the resting diameter of their nuclei, (Davidson *et al*. 2014; Thiam *et al*. 2016; Renkawitz *et al*. 2019). Such experiments generated novel mechanistic insights into this process, including how the nuclear envelope and DNA are repaired after damage caused by severe morphology changes (Raab *et al*. 2016; Denais *et al*. 2016; Xia *et al*. 2018). These approaches mostly rely on migrations through fabrications or physical manipulations of cells in tissue culture and very little is understood about how nuclei traverse confined spaces *in vivo*.

*Caenorhabditis elegans* hypodermal P cells are excellent *in vivo* model for studying nuclear migration through constricted spaces as a normal part of development (Chang *et al*. 2013; Bone *et al*. 2016; Starr 2019). Six P cells are born on each lateral side of the embryo and extend protrusions to form the hypodermis that encloses the ventral surface of the embryo (Williams-Masson *et al*. 1997). P-cell nuclei remain at their original lateral positions throughout embryonic morphogenesis and into the early L1 larval stage when P cells cover the ventral surface of larvae (Hall and Altun 2008). Throughout early L1, the hypodermal hyp7 syncytium intercalates and replaces P cells to cover the ventral surface. During this process, P cells narrow along their anterior-posterior axis and at mid L1, P-cell nuclei migrate from their lateral positions to the ventral cord (Bone *et al*. 2016). P-cell nuclear migration occurs through a constricted space between the body wall muscles and the cuticle where the hypodermis narrows to ∼5% of the resting diameter of a nucleus in early L1. This constriction is small so that the hemidesmosome-like fibrous organelles can form in the hypodermis to mechanically connect body wall muscles to the cuticle (Francis and Waterston 1991; Cox and Hardin 2004; Bone *et al*. 2016). While the fibrous organelles are removed just prior to P-cell nuclear migration, P-cell nuclei still undergo severe morphological changes as they squeeze through this narrow opening (Bone *et al*. 2016). Soon after migrating to the ventral cord, P cells divide and their lineages give rise to the vulval cells and 12 GABA (γ-aminobutyric acid) neurons in the ventral cord (Sulston and Horvitz 1977). Consequently, if P-cell nuclear migration fails, P cells die and the animal develops Egl (<u>eg</u>g-laying deficient) and Unc (<u>unc</u>oordinated) phenotypes due to the absence of vulval cells and neurons, respectively (Sulston and Horvitz 1981; Malone *et al*. 1999; Starr *et al*. 2001).

Multiple molecular pathways contribute to ensure P-cell nuclear migration. The LINC complex pathway consists of UNC-84 and UNC-83, which function together to span the nuclear envelope (Starr and Fridolfsson 2010; Starr 2019). The SUN (<u>S</u>ad1 and <u>UN</u>C-84) protein UNC-84 is an integral protein of the inner nuclear membrane with a N-terminal nucleoplasmic domain that interacts with lamins (Bone *et al*. 2014) and a C-terminal luminal domain that extends across the perinuclear space with its SUN domain that is conserved across eukaryotes being found adjacent to the inner leaflet of the outer nuclear membrane (Cain *et al*. 2014; Jahed *et al*. 2019). The KASH (klarsicht, ANC-1 and Syne Homology) protein UNC-83 is found in the outer nuclear membrane with a short conserved C-terminal KASH peptide of 17 residues that extend into the perinuclear space and a large N-terminal cytoplasmic domain that interacts with microtubule motors (Starr *et al*. 2001; Meyerzon *et al*. 2009; Fridolfsson *et al*. 2010; Fridolfsson and Starr 2010). The KASH peptide of UNC-83 and the SUN domain of UNC-84 directly interact within the perinuclear space to form a physical bridge across the nuclear envelope, connecting the cytoskeleton to the nuclear lamina (McGee *et al*. 2006; Cain *et al*. 2018). During P-cell nuclear migration, the LINC complex works primarily by recruiting the dynein/dynactin complex to the nuclear envelope to move nuclei toward the minus ends of microtubules (Bone *et al*. 2016; Ho *et al*. 2018).

Null mutations in *unc-84* or *unc-83* cause a temperature sensitive phenotype (Malone *et al*. 1999; Starr *et al*. 2001). At 15°C, most P-cell nuclei migrate normally but at 25°C, about 50% of P-cell nuclei fail to migrate (Sulston and Horvitz 1981; Malone *et al*. 1999; Starr *et al*. 2001; Chang *et al*. 2013). This suggests that at least one additional pathway acts in parallel with the LINC complex pathway during P-cell nuclear migration. We screened for additional players in P-cell nuclear migration by mutagenizing and selecting for strains where P-cell nuclear migration failed in an *unc-84* background at the normally permissive temperature of 15 °C (Chang *et al*. 2013). This forward genetic approach identified *toca-1*, which encodes a F-bar protein that localizes to curved membranes and is thought to promote the nucleation of actin filaments through the small GTPase CDC-42 and the Arp2/3 complex (Ho *et al*. 2004; Giuliani *et al*. 2009; Chang *et al*. 2013) and *cgef-1*, a predicted GEF (guanine nucleotide exchange factor) for CDC-42 (Chan and Nance 2013; Ho *et al*. 2023). CDC-42, the Arp2/3 complex member ARX-3, and non-muscle myosin NMY-2 also function in the actin-based pathway (Ho *et al*. 2023).

Here we report that FLN-2 enhances the P-cell nuclear migration defect of *unc-84*. FLN-2 is related to filamins, which usually crosslink or bundle actin filaments (DeMaso *et al*. 2011; Kovacevic *et al*. 2013). However, we showed that the N-terminal actin-binding domain of FLN-2 was not necessary for its function during P-cell nuclear migration through constricted spaces and that the actin network is not severely disrupted in P cells of *fln-2* mutant animals. Instead, we identified a stretch of immunoglobulin (Ig) repeats in FLN-2 that are necessary for P-cell nuclear migration and found an increase in nuclear rupture in *fln-2* mutant P cells. Furthermore, *cgef-1* and *fln-2* had a synergistic nuclear migration defect, suggesting that FLN-2 is a member of a third mechanism that is necessary to move P-cell nuclei in parallel to the LINC complex and actin-based pathways.

## Materials and Methods

### *C. elegans* genetics, strains, and gene editing

Animals were maintained on NGM (nematode growth media) plates spotted with *E. coli* strain OP50 and maintained at 15°C unless otherwise noted (Brenner 1974; Stiernagle 2006). The strains used in this study are listed in Table 1. FX4687 was a gift from Shohei Mitani (Tokyo Women’s Medical University) (Consortium 2012). Some strains were obtained from the Caenorhabditis Genome Center, which is funded by the NIH Office of Research Infrastructure Programs (P40 OD010440).

**Table 1:**
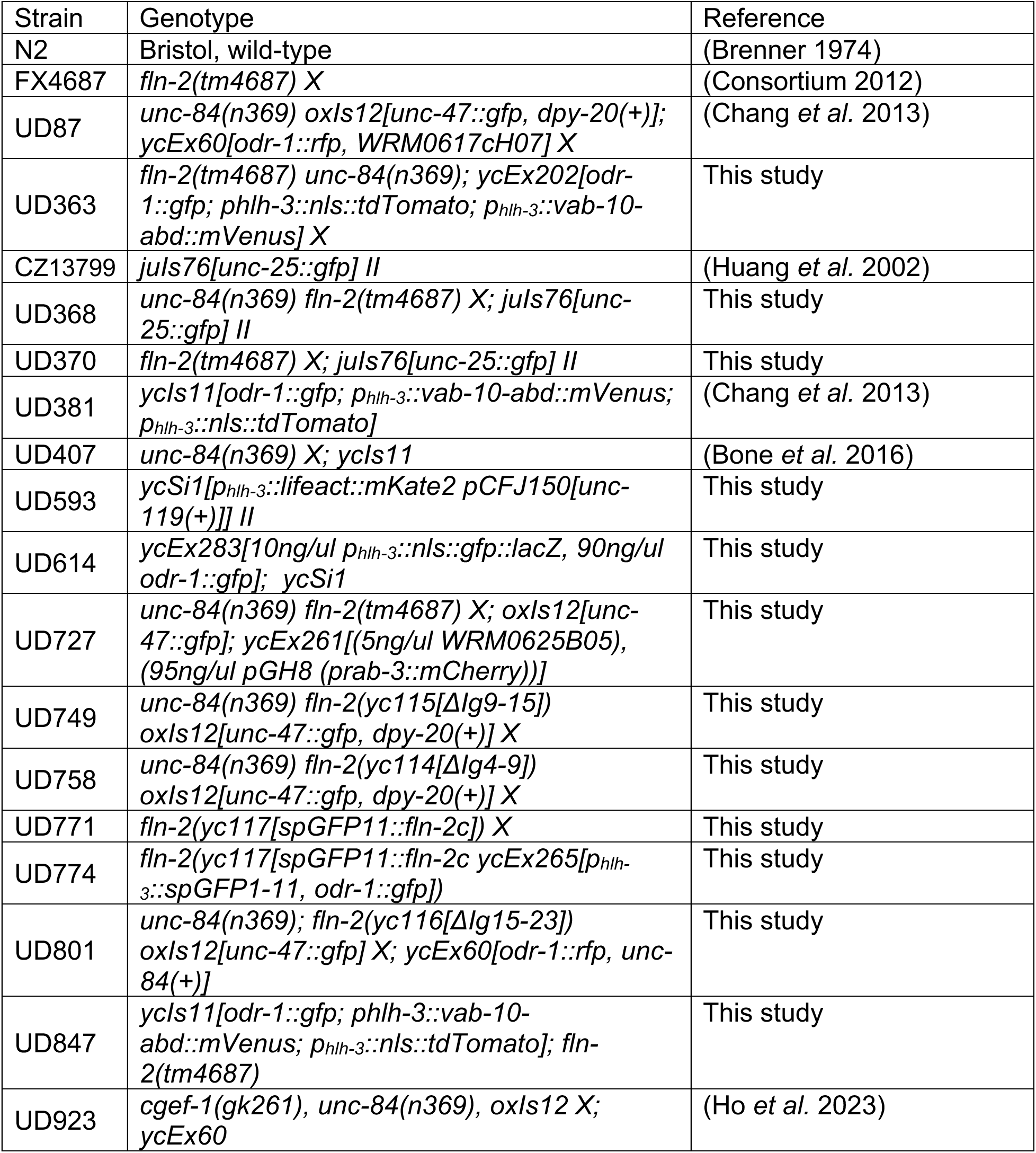
List of worm strains used in this study.

Some strains were made by injecting plasmid DNA to make extrachromosomal arrays (Evans 2006). The plasmid encoding the *vab-10-abd::mVenus* (Kim *et al*. 2011) was cloned under control of the P-cell specific *hlh-3* promoter (Doonan *et al*. 2008; Chang *et al*. 2013). DNA for *phlh-3::nls::tdTomato* was previously described (Chang *et al*. 2013). pSL832 *[phlh-3::nls::gfp lacZ]* was created by amplifying *nls::gfp::LacZ* from pPD96.04 (a gift from Andrew Fire, Addgene plasmid # 1502) and inserting this amplicon into a pSL780 backbone (*phlh-3::lifeact::mKate2)*, replacing the *lifeact::mKate2* with our *nls::gfp::lacZ*. This plasmid was microinjected into UD593 animals to create UD614 (*ycEx283[10ng/ul phlh3::nls::gfp::lacZ, 90ng/ul odr-1::gfp] ycSi1[phlh3::lifeact::mKate2]II)*.

The *fln-2::spGFP11* strain and the *fln-2* domain deletion alleles were produced via CRISPR/Cas-9 gene editing using the crRNA guides and repair templates found in Table 2. We used a *dpy-10* co-CRISPR approach, as previously described (Arribere *et al*. 2014; Paix *et al*. 2017; Hao *et al*. 2021). Two crRNA guides and one single-stranded oligodeoxynucleotide (ssODN) repair template were used to generate each of the *fln-2* deletion alleles (Table 2). The split-GFP approach (Cabantous *et al*. 2005) was used and the eleventh beta-strand of GFP (spGFP11) was inserted to the 5’ end of the predicted open reading frame of *fln-2c*, while the rest of GFP (spGFP1-10) was expressed from an extrachromosomal array under the control of the *hlh-3* promoter (Doonan *et al*. 2008; Chang *et al*. 2013). CRISPR/Cas9 injection mixes were prepared as follows: 0.2 μl of *dpy-10* crRNA (8 μg/ul) (Dharmacon/ Horizon Discovery), a total of 1.5 ul of the target gene crRNA (4μg/ul) (0.75μl of each guide if there are two guides) (Dharmacon/ Horizon Discovery), 1.9μl of tracrRNA (4 μg/μl) (Dharmacon/ Horizon Discovery), 0.5 μl of *dpy-10* ssODN, 1.1μl of the target gene ssODN (1 μg/μl), 4 μl of Cas9 (10 μg/μl) (UC Berkeley QB3), and 0.8 μl of H2O. The mix was injected into the gonads of young adult hermaphrodite animals (Evans 2006). All ssODN repair templates were manufactured by Integrated DNA Technologies.

**Table 2.**
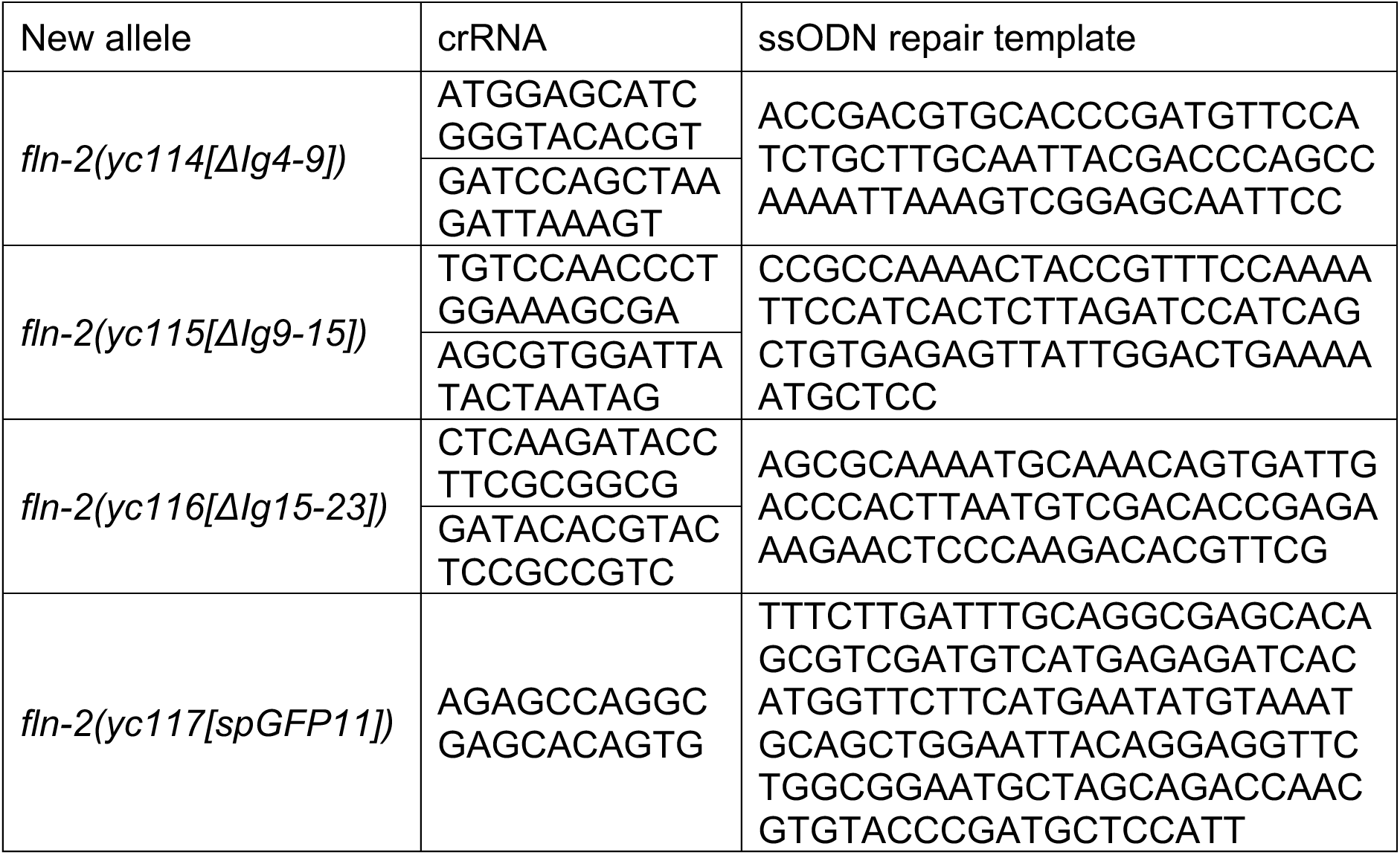
Sequences of CRISPR crRNA guides and ssODN repair templates used in this study.

The lifeact::mKate2 in P cell strain (UD593) was made using *mos1*-mediated single copy insertion (mosSCI) (Frøkjær-Jensen *et al*. 2014). DNA encoding the *hlh-3* promoter (Doonan *et al*. 2008), lifeact (Riedl *et al*. 2008), and mKate2 (Shemiakina *et al*. 2012) were from pSL780 (Bone *et al*. 2016) and cloned into Gateway entry plasmids (pDONR 221 P4-P1R, pDONR 221, and pDONR 221 P2r-P39 (ThermoFisher Scientific)). Subsequently, they were cloned using the Gateway LR reaction into the pCFJ150 targeting vector (a gift from Erik Jorgensen; Addgene plasmid # 19329) (Frøkjær-Jensen *et al*. 2008) and named pSL831. This plasmid was injected into EG6699 (*ttTi5605)* animals (Frøkjær-Jensen *et al*. 2012) along with pCFJ601 *(Peft-3* Mos1 transposase; 50 ng/μl), pMA122 (*peel-1* negative selection; 10 ng/μl), pGH8 (*Prab-3::mCherry;* 10ng/ul), pCFJ90 (*Pmyo-3::twk-18(cn110)*; 2.5ng/μl), and pCFJ104 *(Pmyo-3::mCherry)* (5 ng/μl). Animals harboring a single copy of integrated pSL381 were selected as *unc-119(+)* and lacked the red extrachromosomal array markers (Frøkjær-Jensen *et al*. 2008, 2012)

### *C. elegans* P-cell nuclear migration assays, synchronization, and microscopy

Our assay for counting failed P-cell nuclear migration events using *unc-47::gfp* to mark GABA neurons in the ventral cord has been previously described (Chang *et al*. 2013; Fridolfsson *et al*. 2018)

For some experiments, *C. elegans* animals were roughly synchronized in the middle of the L1 larval stage when P-cell nuclear migration takes place (Sulston and Horvitz 1977). Six plates of gravid hermaphrodites were bleached as described (Stiernagle 2006) and the surviving embryos were split between three plates to develop at 15° C, 18° C, or 20° C for 18 hours. The plates with the most L1 animals in the middle of P-cell nuclear migration were identified by examining P-cell nuclei with the *phlh-3::nls::tdTomato* marker using a Leica FLIII fluorescent dissecting stereo microscope. L1 animals were washed off the plate with M9 buffer (Stiernagle 2006) and mounted on a 2% agarose pad. Then, tetramizole was added to a final concentration of 1 μM as a paralytic, and a cover slip was applied and sealed with VALAP (equal parts Vaseline, lanolin, and paraffin mixture). L1 larvae were imaged with the Zeiss LSM 980-Airyscan2 with a 63x NA 1.4 objective in the MCB Light Microscopy Imaging Facility at UC Davis made available through an NIH grant S10OD026702.

### Alphafold

Alphafold models of FLN-1 and FLN-2 were generated using ColabFold version 1.5.2 (Mirdita *et al*. 2022) using the ‘alphafold2_ptm’ model. No templates were used, and 5 models were generated (using default settings). The models shown have the highest pLDDT (predicted local distance difference test) score. Protein diagrams were colored by sequence order from blue (N terminus) to red (C terminus) using the ‘rainbow’ command in UCSF ChimeraX, developed by the Resource for Biocomputing, Visualization, and Informatics at UCSF with support from National Institutes of Health R01-GM129325 and the Office of Cyber Infrastructure and Computational Biology, National Institute of Allergy and Infectious Diseases (Pettersen *et al*. 2021). To disentangle the folded Ig domains from disordered loops, low-pLDDT regions were stretched out by assigning each torsion angle drawn from a normal distribution (phi=-165° +/- 5°, psi=165 +/- 5°, where 5° is the standard deviation).

## Results

### *fln-2* is an enhancer of the nuclear migration defect observed in *unc-84* animals

A previously performed screen was used to identify enhancers of the nuclear migration defect of *unc-84* (i.e., *emu* genes), which function parallel to LINC complexes during P-cell nuclear migration (Chang *et al*. 2013). While analyzing mutant alleles from an *emu* screen, we found that mutants in *fln-2* enhance the nuclear migration defect of *unc-84* (Figures 1A-D). We counted the number of GABA neurons present in young adults using an established *unc-47::gfp* marker (McIntire *et al*. 1997). Since 12 of the 19 GABA neurons are derived from P-cells (Sulston and Horvitz 1977), the absence of GABA neurons can be used as a proxy for scoring the number of P-cells that died during nuclear migration in L1 larvae (Figure 1A-C) (Chang *et al*. 2013; Fridolfsson *et al*. 2018). Thus, a completely penetrant P-cell nuclear migration defect would result in 12 missing GABA neurons. Animals harboring *fln-2(tm4687),* an expected null allele, had a mild P-cell nuclear migration defect with an average of 2.4 missing GABA neurons at 15°C (Figures 1A-D). Both *fln-2(tm4867)* and *fln-2(RNAi)* animals significantly increased the nuclear migration defects observed in *unc-84* mutant animals with an average of 6.6 and 4.6 missing GABA neurons, respectively in the double mutants (Figure 1D). Therefore, *fln-2* likely plays a role during P-cell nuclear migration.

**Figure 1:**
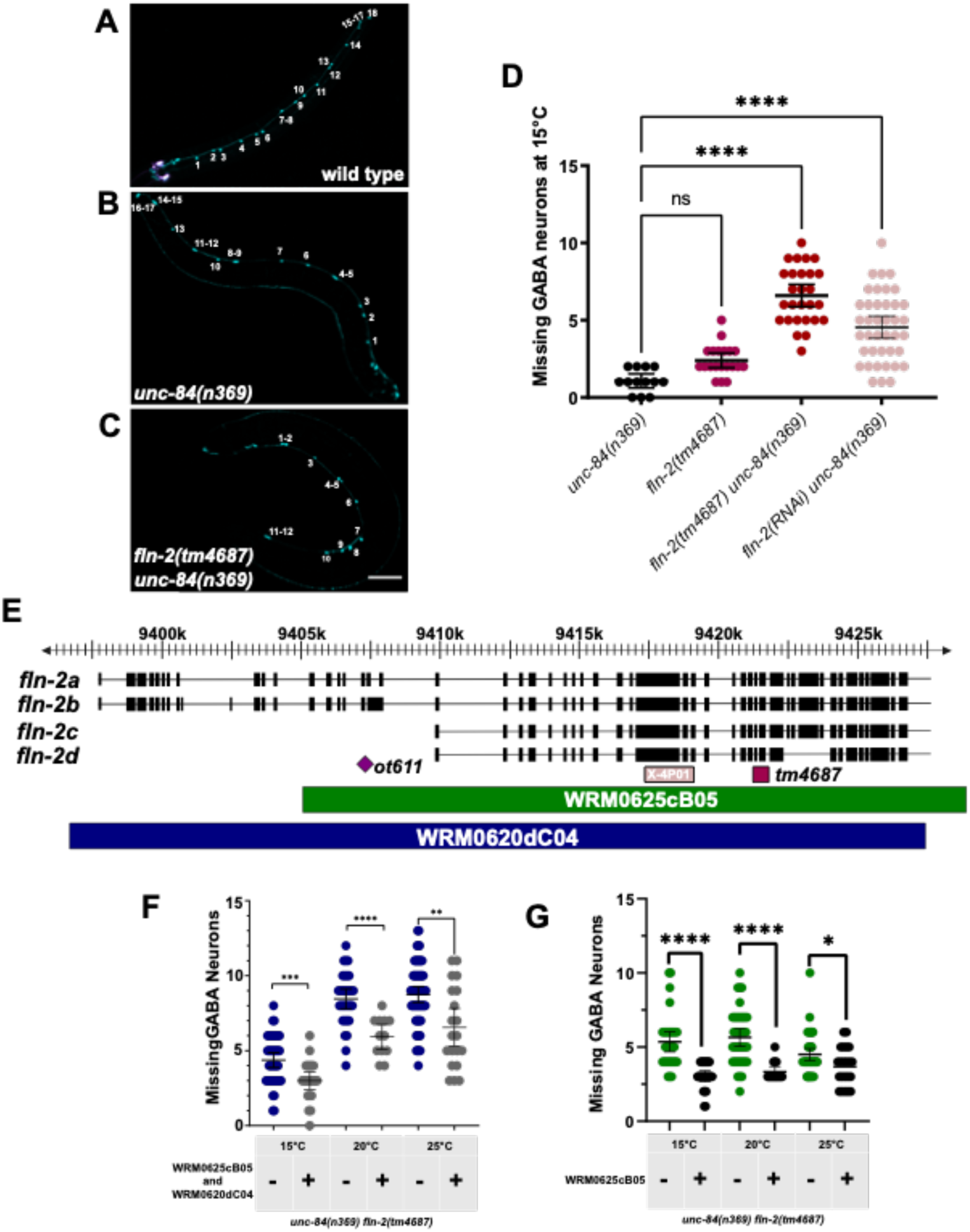
*fln-2* mutants enhance the P-cell nuclear migration defect observed in *unc-84* worms. (A-C) Representative epifluorescence images of *unc-47::gfp-*marked GABA neurons in wild type, *unc-84(n369)*, or *unc-84(n369); fln-2(tm4687)* animals. (D) The numbers of missing GABA neurons are shown from the following genotypes: *unc-84(n369)* (black), *fln-2(tm4687)* (magenta), *unc-84(n369) fln-2(tm4687)* (red), and *unc-84(n369) fln-2(RNAi)* (pink). **** signifies a P-value < 0.0001 when compared to *unc-84(n369)* alone. (E) Schematic of four of the more than 25 predicted isoforms of *fln-2*. The purple diamond indicates the *ot611* point mutant producing an early stop codon. The pink box indicates the X-4P01 clone used for RNAi. The magenta box shows the location of the *tm4687* deletion allele. The fosmids used in these experiments are marked below the isoforms. (F-G) Missing GABA neuron counts for the *unc-84(n369) fln-2(tm4687)* mutants with and without the indicated fosmids. *, **, ***, and **** indicates a p-value < 0.02, < 0.003, 0.0007, and < 0.0001, respectively; error bars are 95% confidence intervals.

According to Wormbase (www.wormbase.org), over 25 *fln-2* isoforms are predicted to exist, four of which are presented in Figure 1E (DeMaso *et al*. 2011). We aimed to determine which *fln-2* isoforms were necessary for P-cell nuclear migration. The *fln-2(tm4687)* allele, which has a 547 bp deletion that is predicted to cause a frameshift and disrupt most *fln-2* isoforms, caused a strong nuclear migration defect in the *unc-84(null)* background (Figure 1E). Furthermore, *fln-2(RNAi)* performed using clone X-4P01 (Fraser *et al*. 2000; Kamath *et al*. 2003) caused a similar phenotype. In contrast, the *fln-2(ot611)* mutant, which introduces an early stop codon in the longer *fln-2* isoforms, was found in our background strain, UD87, that was used for our enhancer screen (Chang *et al*. 2013). Therefore, the *ot611* allele has no phenotype, suggesting that the longer *fln-2* isoforms are dispensable for P-cell nuclear migration (Figures 1A-E). To confirm that the *fln-2(tm4687)* P-cell nuclear migration defect was caused by the deletion in *fln-2* and not somewhere else in the genome of this strain, we performed fosmid rescue experiments (Figures 1E-G). A combination of two fosmids (i.e., WRM0625cB05 and WRM0620dC04) that span the entire *fln-2* locus rescued the *fln-2(tm4687) unc-84(n369)* P-cell nuclear migration defect (Figure 1F). Furthermore, a shorter fosmid, WRM0625cB05, that did not cover the longer *fln-2* isoforms was also able to rescue *fln-2(tm4687) unc-84(n369)* (Figure 1G). These results support the hypothesis that the smaller isoforms *fln-2c* or *fln-2d* are sufficient for P-cell nuclear migration.

### Ig-like repeats 4-8 of FLN-2 are necessary for P-cell nuclear migration

Like other filamin family members, the shorter isoforms *fln-2c/d* are predicted to encode proteins mostly made of tandem Ig-like repeats (DeMaso *et al*. 2011). We performed CRISPR/Cas9-mediated gene editing to generate in-frame deletions in the endogenous *fln-2* locus to better determine which FLN-2 Ig-like repeats were necessary for P-cell nuclear migration. *fln-2c* consists of 2776 amino acids and has 20 predicted Ig-like repeats (Figure 2A). We divided FLN-2c into three overlapping segments, Ig repeats 4-9, 9-15, or 15-23 (DeMaso *et al*. 2011) and made three different deletion strains (Figure 2A). Each deletion strain was analyzed for defects in P-cell nuclear migration by counting the number of missing GABA neurons present at 15°C in the presence or absence of *unc-84* (Figure 2). Deletion of Ig-like repeats 4-9 significantly enhanced the *unc-84(null)* nuclear migration defect (Figure 2B), whereas deletion of repeats 9-15 or 15-23 had no effect (Figure 2C-D). These results narrow down a part of FLN-2 that is necessary for P-cell nuclear migration to Ig-like repeats 4-8, which includes the disordered region between repeats 6 and 7.

**Figure 2:**
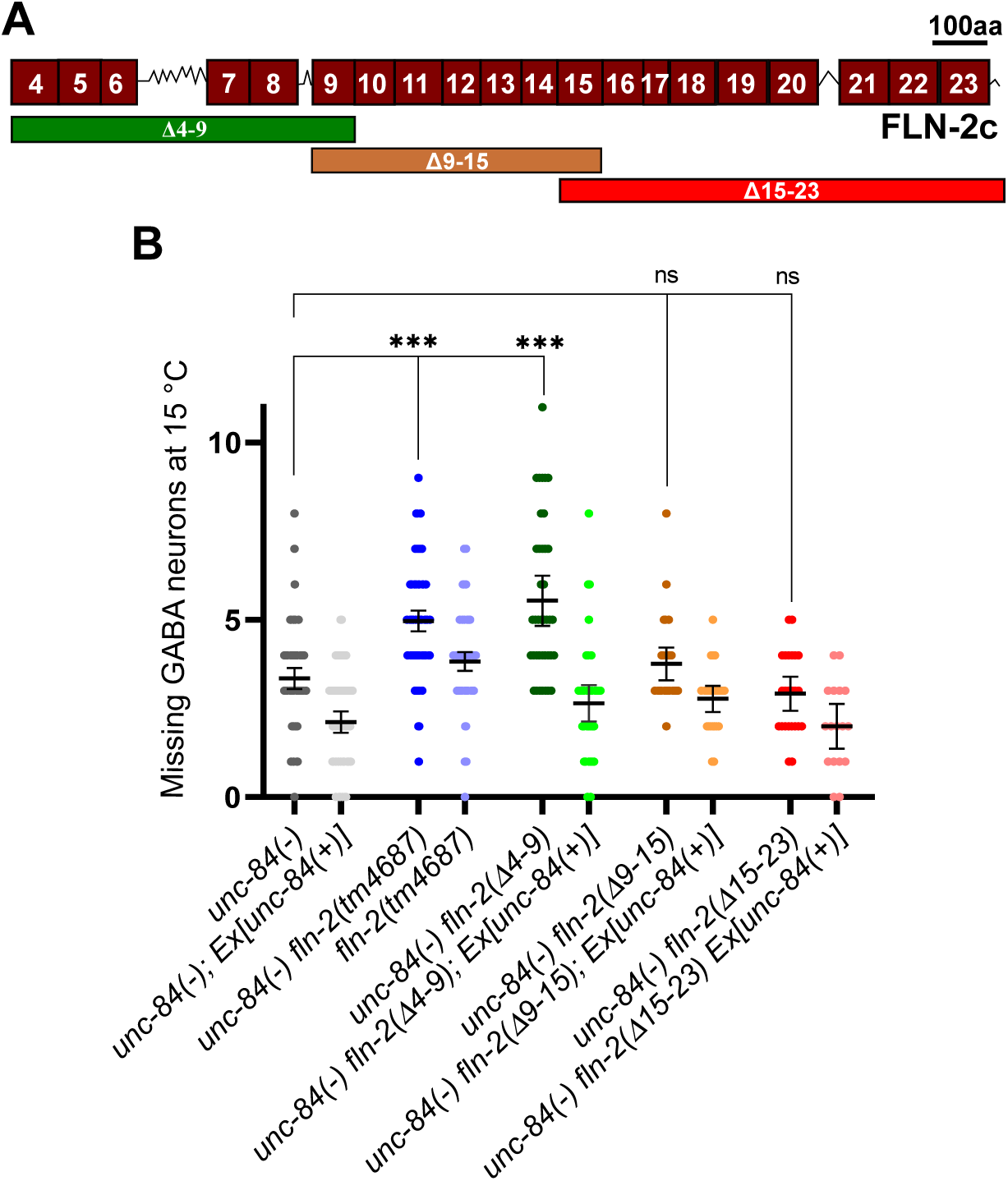
Deletion of Ig-like repeats 4-9 enhances the *unc-84* P-cell nuclear migration defect. (A) Schematic of the FLN-2c isoform, which contains 20 predicted Ig-like repeats (maroon boxes). Ig-like repeats 1-3 are only found in the longer isoforms of FLMN-2 with interspersed disordered regions are indicated by the zig-zag lines. The region between repeats 6 and 7 is predicted to be disordered. The deleted portions of FLN-2 in three strains made by CRISPR/Cas9 editing are indicated below the schematic. The scale bar indicates 100 amino acids. (B) Missing GABA neurons at 15°C were scored as a proxy for failed P-cell nuclear migrations. See Table 1 for details on the genotypes of the strains used here. Means with 95% confidence intervals are shown. *** indicates significant differences with a p-value of <0.001 when compared to *unc-84* null animals and ns stands for no significant difference from *unc-84* null animals. A t-test was done to generate p-values.

### The actin network in P cells is not grossly disrupted in *fln-2* mutants

Since filamins are thought to cross link and bundle actin (Razinia *et al*. 2012) we examined the organization of the actin cytoskeleton during P-cell nuclear migration in *fln-2* mutant L1 larvae. We compared two reporters for actin filaments—the small actin-binding domain from *vab-10* bound to a fluorescent protein (*vab-10::mVenus*) under control of the P-cell-specific *hlh-3* promoter expressed from a high-copy number extrachromosomal array (Bone *et al*. 2016) and a single-copy transgene expressing *lifeact::mKate2* (Riedl *et al*. 2008) under the *hlh-3* promoter inserted into a known locus (*ttTi5605*) using *mosSCI* (Frøkjær-Jensen *et al*. 2014). Both the over-expressed *vab-10::mVenus* (Figure 3) and the single-copy *lifeact::mKate2* markers localized in similar patterns when imaged with Airyscan super-resolution microscopy.

**Figure 3:**
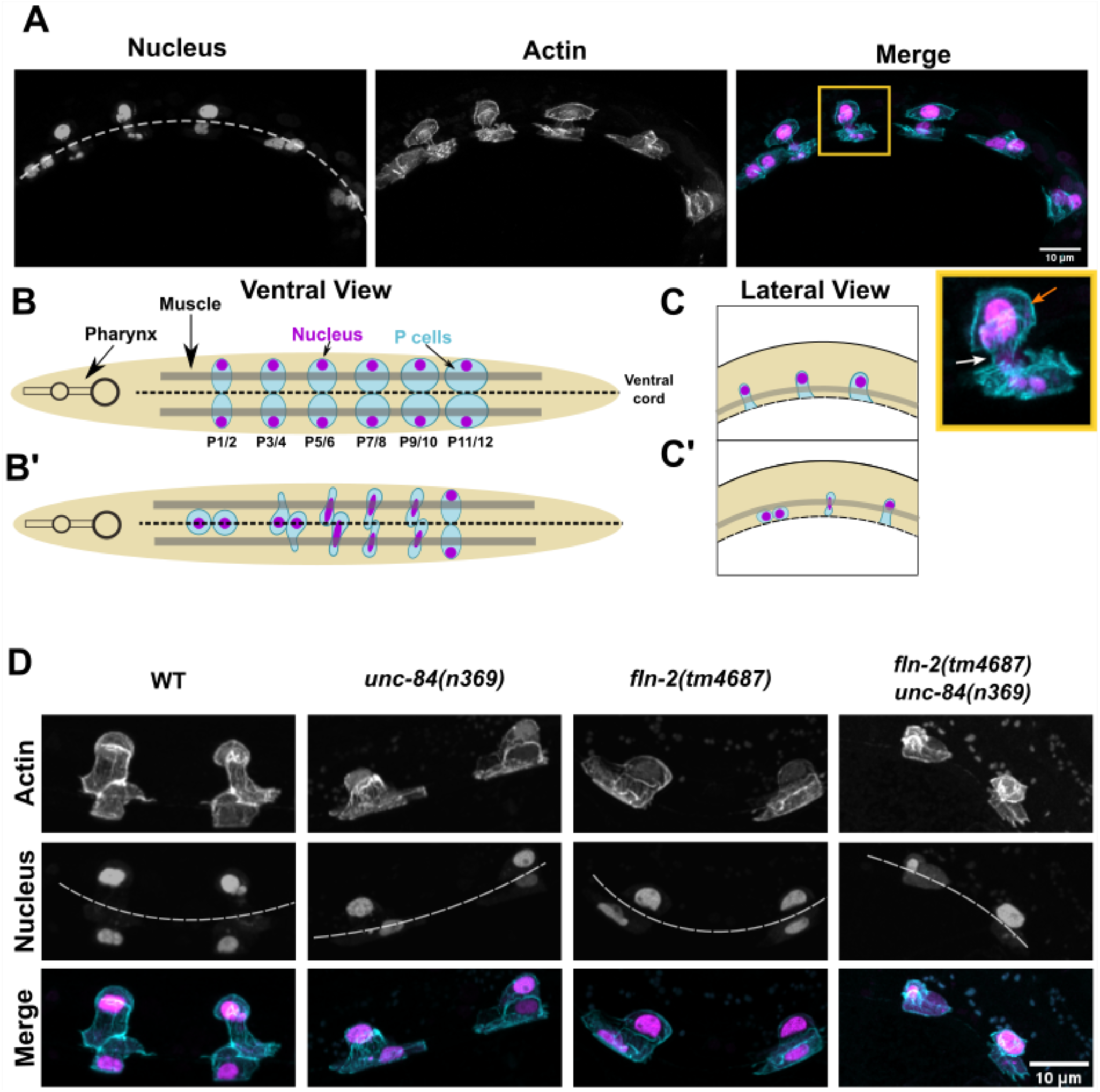
Actin networks during P-cell nuclear migration. (A) Representative Airyscan super-resolution images of a wild-type L1 larvae during P-cell nuclear migration. P-cell nuclei are marked by *nls::tdTomato* (magenta) and actin in P cells is tagged with *vab-10::mVenus* (cyan). The gold box indicates a P cell mid nuclear migration that is enlarged in the inset below. The orange arrow points out actin perpendicular to the direction of nuclear migration in the lateral compartment and the white arrow indicates actin parallel to the direction of nuclear migration in the constriction. A ventral view is shown and the ventral cord is marked by the dashed line. Anterior is left; scale bar is 10 μm. (B-C) A ventral view (B) and lateral view (C) of P cells during the narrowing (B and C) and migration (B’ and C’) stages. P cells are in cyan, nuclei are in magenta. The dashed line indicates the ventral cord. The pharynx indicates the anterior end of the worm and body wall muscles are represented by the transparent gray bar. (D) Actin (*vab10::mVenus*; cyan) and P-cell nuclei (*nls::tdTomato*; magenta) in wild-type, *unc-84(n369)*, *fln-2(tm4687)*, or *fln-2(tm4687) unc-84(n369)* double mutant animals. Maximum projections of z stacks are shown. The scale bar is 10 μm.

First, we characterized the wild-type organization of actin filaments during the narrowing and migration stages of P-cell nuclear migration (Bone *et al*. 2016). The most anterior pair of P cells narrow, and their nuclei migrate earlier than the more posterior pairs. During nuclear migration, pairs of P-cell nuclei deform and move through the narrow constriction between the cuticle and body wall muscles to reach the ventral cord (Bone *et al*. 2016). Actin filaments were present both behind the nucleus and in the constriction during P-cell nuclear migration in 16 out of 16 worms (Figure 3A). We frequently observed actin bundles forming perpendicular to the direction of nuclear migration in the lateral portion of P-cells behind migrating nuclei (orange arrows in Figure 3A). Additionally, we observed actin fibers forming parallel to the direction of nuclear migration within the constriction (white arrows in Figure 3A). We then compared the organization of actin networks in our mutant animals. In *unc-84(n369)* or *fln-2(tm4687)* single mutants, and in *fln-2(tm4687) unc-84(n369)* double mutants, the actin networks did not have any gross changes as compared to wild type (Figure 3D), suggesting that *fln-2* is not required for the presence or organization of actin networks during P-cell nuclear migration.

### *fln-2* mutant P-cell nuclei rupture post-migration

The nuclei of cultured mammalian cells migrating through fabricated constrictions often rupture, releasing soluble nucleoplasmic proteins into the cytoplasm (Raab *et al*. 2016; Denais *et al*. 2016). We examined the extent to which P-cell nuclei rupture during migration though constricted spaces in *C. elegans* L1 larvae. We followed nuclear rupturing with the localization of a soluble nuclear marker (nls::tDtomato) to see if it leaked into the cytoplasm before, during, or after P-cell nuclear migration. Normally, nls::tDtomato was restrained within the nucleus, but after nuclear rupturing, the fluorescent signal was dilute and detected throughout the cytoplasm in both the lateral and ventral compartments of P cells (arrowheads in Figure 4A). In wild type L1 larvae, there were rarely any ruptured nuclei prior to or during nuclear migration. However, a few wild type nuclei ruptured toward the end of the nuclear migration process (Figure 4B). The number of nuclear rupture events observed in late migration was significantly increased in both *unc-84(n369)* and *fln-2(tm4687)* single mutant larvae relative to wild type. Furthermore, *fln-2(tm4687), unc-84(n369)* double mutant animals had a significant enhancement of the nuclear rupture phenotype (Figure 4B). These data suggest that *unc-84* and *fln-2* function synergistically in maintaining the integrity of the nuclear envelope during P-cell nuclear migration.

**Figure 4:**
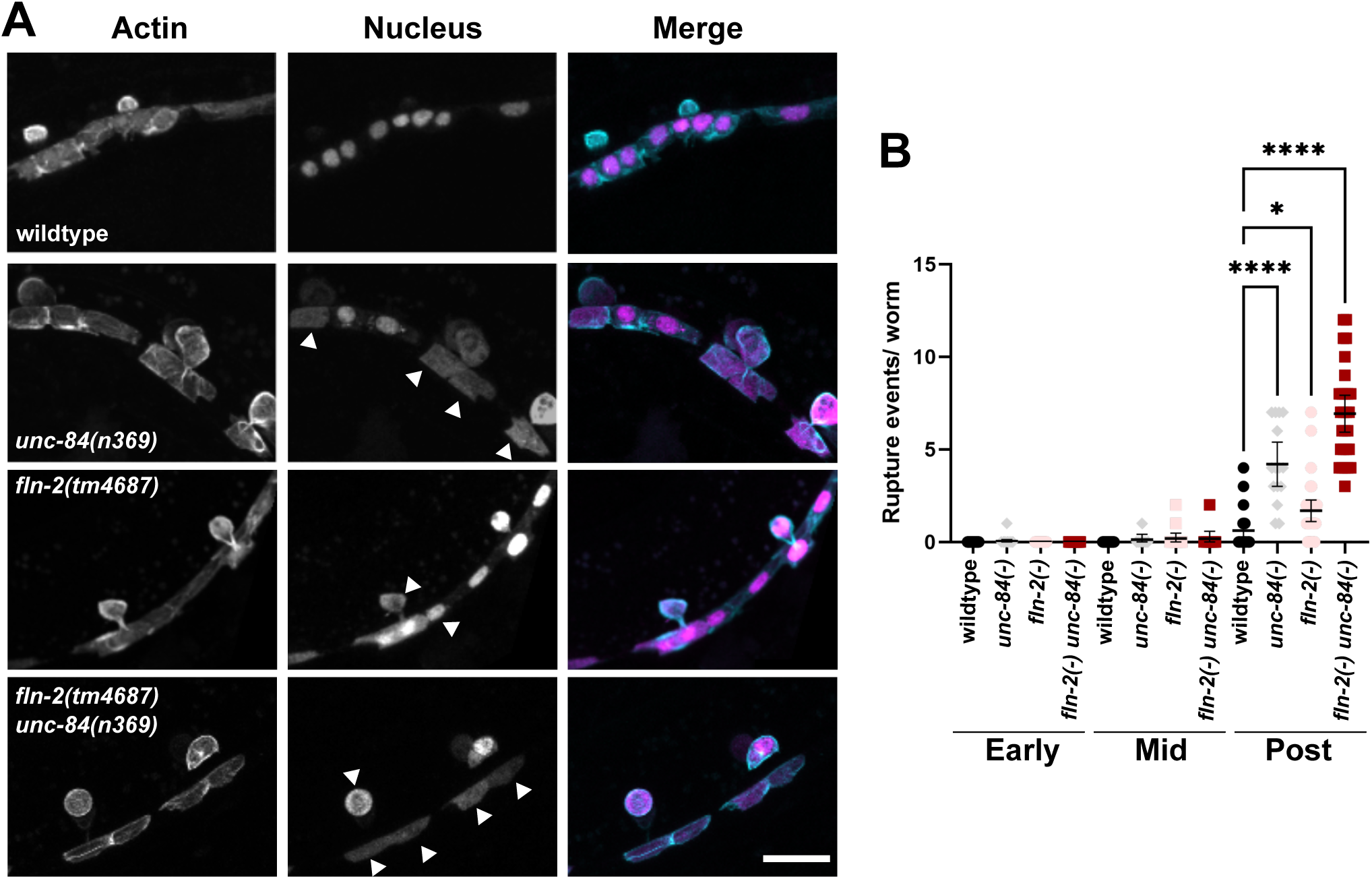
P-cell nuclei in *fln-2 unc-84* mutant animals often rupture during migration. (A) Representative Airyscan super-resolution images of P cells late in L1 after or during nuclear migration into the ventral cord in wildtype, *unc-84(n369)*, *fln-2(tm4687)*, and *fln-2(tm4687) unc-84(n369)* animals. Actin is shown to mark the shape of P cells. Arrows point to suspected nuclear rupture events where the nls::tdTomato marker has leaked into the cytoplasm of the P cell. Maximum projections of z stacks are shown. The scale bar is 10 μm. (B) The number of P cells with cytoplasmic nls::tdTomato per worm in late L1. Means and 95% confidence intervals are shown. **** denotes a p-value of <0.00001.

### FLN-2 localization in *C. elegans* larvae

Canonical filamins directly interact with actin (Razinia *et al*. 2012). We therefore explored the extent to which FLN-2 might colocalize with actin networks. We localized FLN-2c *in vivo* using a split-GFP assay where the beta strand 11 of GFP was fused to the endogenous locus for FLN-2c and strands 1-10 of GFP were expressed under the control of a P-cell specific promoter, *hlh-3* (Figure 5A). The spGFP11 tag did not disrupt FLN-2 function; *unc-84(n369); gfp11::fln-2c* animals had no significant defect in the number of GABA neurons when compared to *unc-84(n369)* animals (Figure 5B). spGFP11::FLN-2c localized diffusely throughout the cytoplasm and was slightly enriched in the nucleoplasm (Figure 5A). These experiments suggest that the bulk of FLN-2 does not co-localize to actin networks in P cells.

**Figure 5:**
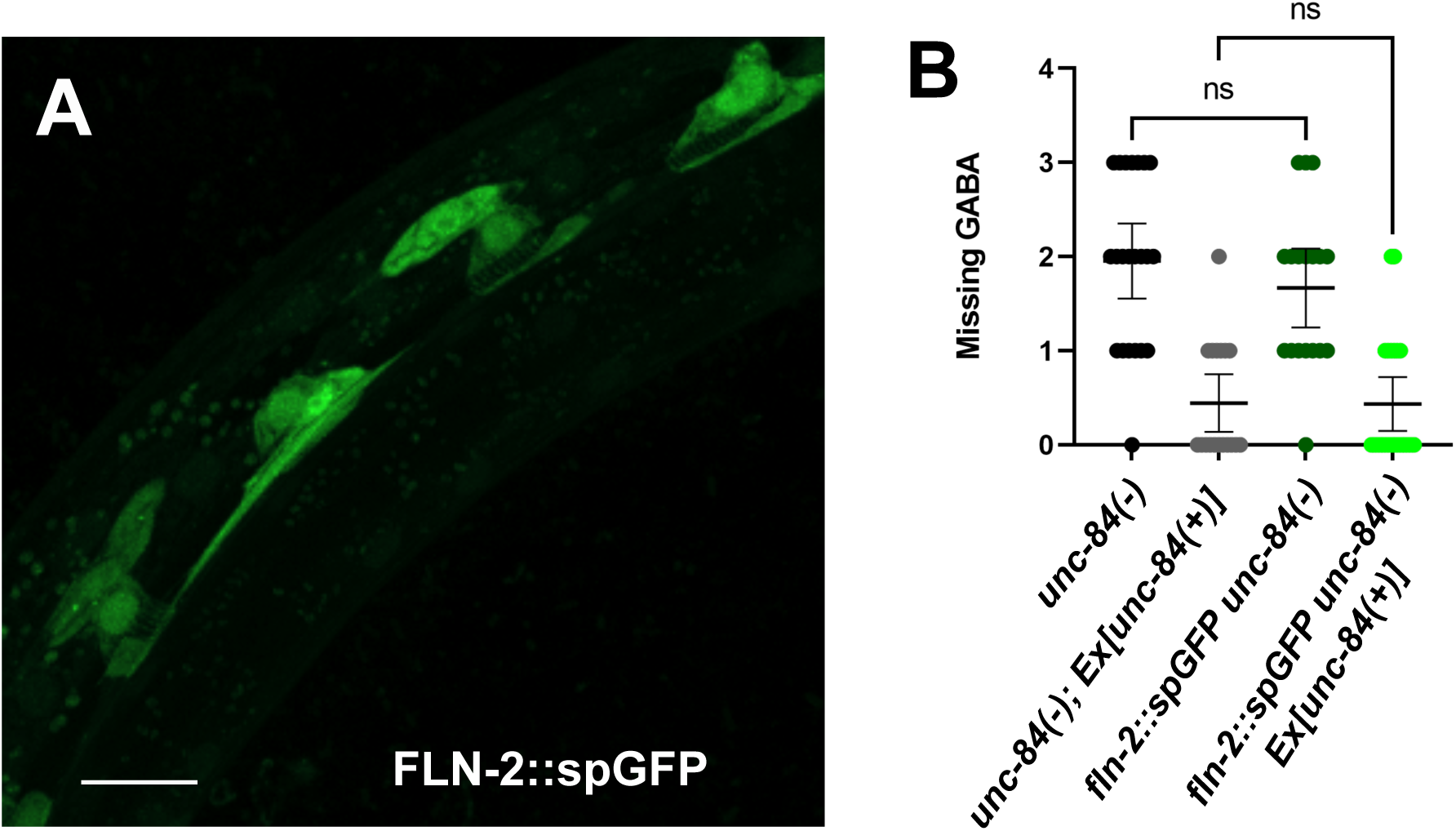
FLN-2 localization in larval P cells. **A)** Representative Airyscan super-resolution image of *fln-2c::GFP* expressed in a *C. elegans* L1 larva. A lateral view is shown; ventral is toward the lower right. Scale bar is 10 μm. **B)** Graph depicting the number of missing GABA neurons in *unc-84(n369)*, *unc-84(n369) + Ex[unc-84(+)]*, *fln-2c::spGFP unc-84(n369)*, and *fln-2c::spGFP unc-84(n369) + Ex[unc-84(+)]* worms. Error bars are 95% CI, and non-significant P-values > 0.05.

### *fln-2* functions in parallel to the LINC complex and actin-based pathways

The *fln-2, unc-84* double mutant phenotype is not completely penetrant. Even at the restrictive temperature, many P-cell nuclei still complete their migration, suggesting that other pathways are still functioning. Double mutants in the CDC-42/actin-based pathway and the LINC complex pathway have a similar, incompletely penetrant, phenotype (Ho *et al*. 2023). We therefore tested whether *fln-2* is in the same actin-based pathway as the *cdc-42* GEF *cgef-1*. To test this hypothesis, we examined P-cell nuclear migration in *fln-2(RNAi), cgef-1(gk261), unc-84(n369)* triple mutant animals. Animals carrying all three mutations had significantly more missing GABA neurons than either *fln-2(RNAi), unc-84* or *cgef-1, unc-84* double mutants (Figure 6). Thus, *fln-2* and *cgef-1* have a synergistic relationship during nuclear migration through confined spaces in P cells. These data are consistent with a model where *fln-2*, *cgef-1*, and *unc-84* act in three parallel pathways to ensure P-cell nuclear migration through constricted spaces.

**Figure 6:**
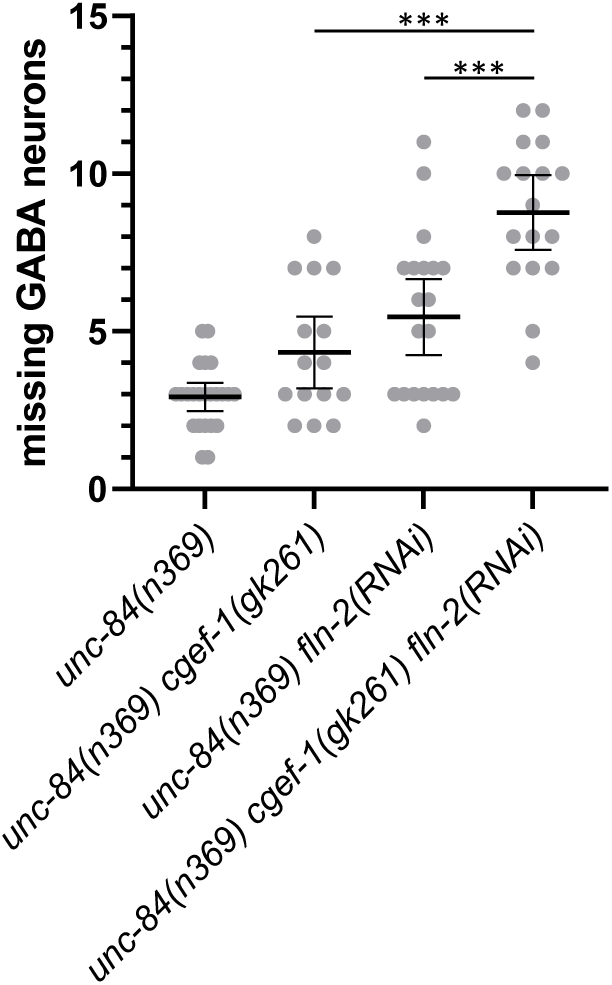
*fln-2* has synthetic interactions with *unc-84* and *cgef-1*. Graph depicting the number of missing GABA neurons in the designated genotypes. The mean is shown with 95% CI error bars. Data was collected at 15°C. n is at least 15. *** denotes *p*<0.001 from a paired t-test.

### FLN-2 might not be a filamin

The above-presented data are not consistent with FLN-2 functioning as a canonical filamin. FLN-2 is quite divergent from canonical filamins, falling far outside of a phylogenetic tree where human FLNA, FLNB, FLNC, *C. elegans* FLN-1, and *Drosophila* Jbug cluster together (DeMaso *et al*. 2011), the CH domain of FLN-2 is not necessary (Figure 1), and the localization of FLN-2::GFP does not suggest it interacts with actin (compare Figures 3 and 5). We therefore used Alphafold to predict the structures of FLN-2d and the canonical filamin FLN-1 (Figure 7). FLN-1 was predicted with high confidence to consist of two N-terminal CH domains (gray in Figure 7) followed by a chain of Ig repeats, as expected. In contrast, only a few Ig repeats were predicted in FLN-2, and instead it was predicted to have extensive disordered stretches.

**Figure 7:**
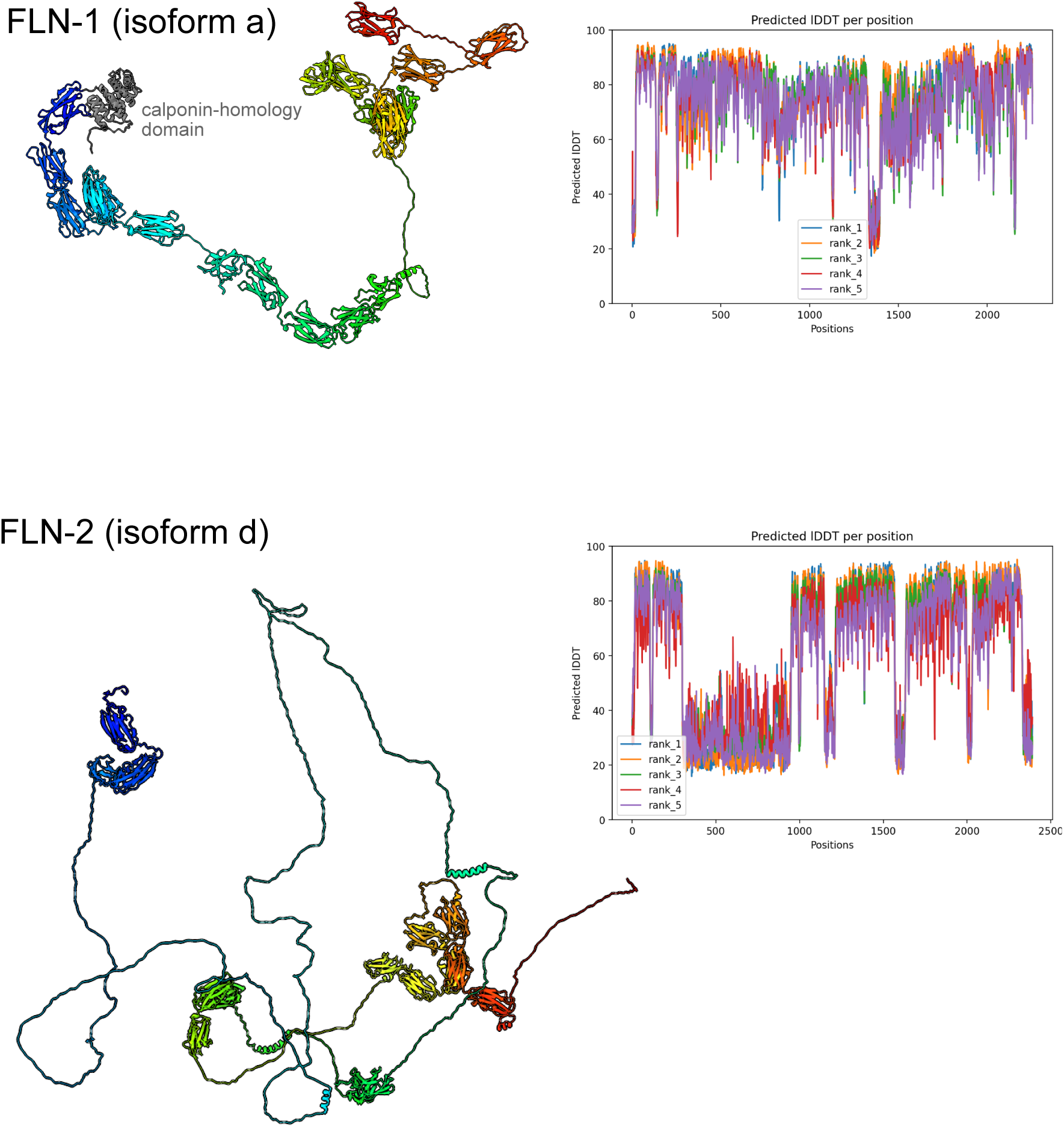
Alphafold predictions of the structure of FLN-1 and FLN-2. The highest confidence of five FLN-1a (top) and FLN-2d (bottom) structures predicted by ColabFold version 1.5.2 are shown. On the right, the pLDDT scores (predicted local distance difference test; a per-residue confidence metric) for five different predictions are shown. The protein diagrams are colored by sequence order from blue (N terminus) to red (C terminus).

## Discussion

We identified *fln-2* as part of a new genetic pathway that functions to move P-cell nuclei through constricted spaces in developing *C. elegans*. Previous studies focused on two other pathways. First, the LINC complex pathway recruits the microtubule motor dynein to the surface of nuclei to move them toward the minus ends of microtubules (Fridolfsson *et al*. 2010; Bone *et al*. 2016; Ho *et al*. 2018). The second pathway is an actin-based process that includes the small GTPase CDC-42, its GEF CGEF-1, TOCA-1, and the Arp2/3 nucleating complex (Chang *et al*. 2013; Ho *et al*. 2023). Our genetic analyses showed that mutations in *fln-2* synergistically interacted with both *unc-84* of the LINC complex pathway and *cgef-1* of the actin-based pathway. Thus, there are at least three genetic pathways that function together to move P-cell nuclei. Since P-cell nuclei migrate as a normal part of development through narrow openings (Bone *et al*. 2016), this system is an excellent model for better understanding how nuclei might migrate through constricted spaces as part of neuronal development, hematopoiesis, inflammation, and metastasis.

One potential model for how FLN-2 could function during nuclear migration through confined spaces in P cells is by organizing the actin cytoskeleton. FLN-2 is a predicted divergent filamin (DeMaso *et al*. 2011), and mammalian filamins function, at least in part, by crosslinking or bundling actin filaments (Nakamura *et al*. 2011). However, our data do not support such a role for FLN-2. First, a short isoform, FLN-2c, that is missing the three N-terminal calponin homology repeats previously predicted to be an actin-binding domain, is sufficient for nuclear migration in P cells. Second, the *fln-2(ot611)* allele, which is predicted to introduce an early stop codon in the longest isoform of FLN-2, is found in many wildtype strains, including our lab’s version of N2, suggesting that the longest isoforms with the predicted actin-binding domains are not selected for any significant functions (Sarin *et al*. 2010). Furthermore, FLN-1 is the canonical filamin in *C. elegans*, while FLN-2 is evolutionarily quite distant from other animal filamins (DeMaso *et al*. 2011), suggesting that it may play divergent functions. Furthermore, Alphafold predictions of FLN-2 did not support the hypothesis that it folds like a canonical filamin, predicting instead that FLN-2 consists of extensive stretches of disordered domains. Third, *fln-2* functions parallel to the CDC-42/actin-based pathway. Finally, *fln-2* mutant animals had grossly normal actin networks in larval P cells. Together, these data do not support a model where FLN-2 functions in the organization of actin networks during P-cell nuclear migration and that it probably is better if FLN-2 is not thought of as a filamin.

An alternative model that we favor is that FLN-2 helps to maintain and/or repair the mechanical integrity of the nuclear envelope during P-cell nuclear migration. *fln-2, unc-84* mutant P-cell nuclei had a significant increase in rupture frequency of the nuclear envelope compared to the single mutants or wild type. Nuclear envelope rupture and subsequent DNA damage regularly occurs when dendritic or cancer cells migrate through fabricated constrictions (Raab *et al*. 2016; Denais *et al*. 2016). Nuclear envelope rupture is repaired by the ESCRT (endosomal sorting complex require for transport) complex (Olmos 2022). Since P cells divide soon after nuclear migration to the ventral cord (Sulston and Horvitz 1977; Hall and Altun 2008), the FLN-2 pathway might function to prevent nuclear rupture and therefore prevent DNA damage during a window when the cell is preparing to divide.

FLN-2 also functions in the formation of multi-vesicular bodies. Three alleles in *fln-2* were found in a forward genetic screen for diminished VHA-5::RFP-containing puncta, a marker for multi-vesicular bodies (Shi *et al*. 2022). *fln-2* mutants also have a reduced number of multi-vesicular bodies as assayed by electron microscopy (Shi *et al*. 2022). Interestingly, both multi-vesicular body formation and nuclear envelope repair have a common molecular mechanism through the ESCRT pathway (Olmos 2022). However, FLN-2 acts at a different step than the ESCRT pathway to regulate multi-vesicular bodies (Shi *et al*. 2022). Shi *et al* (2022) conclude that the N-terminus of the longest FLN-2 isoforms functions by binding both actin filaments and the VHA-8 subunit of multi-vesicular bodies. However, the presence of the *fln-2(ot611)* allele in many wild type strains predicts that the N-terminal domain is dispensable for P-cell nuclear migration. Together, these data suggest a model where shorter isoforms of FLN-2 function in nuclear envelope and multi-vesicular body maintenance and/or repair. The molecular mechanism underlying how FLN-2 functions and interplays with the ESCRT pathway requires further studies.

## Acknowledgements

We thank past and present members of the Starr-Luxton lab for insightful discussions. We thank the Caenorhabditis Genetics Center and WormBase. We thank Thomas Wilkop and the MCB Light Microscopy Imaging Facility at UC Davis. This research was funded by a grant from the National Institutes of Health, R35GM134859 to D.A.S. and the training grant T32GM00737 supported L.M.

## References

1. Arribere J. A., R. T. Bell, B. X. H. Fu, K. L. Artiles, P. S. Hartman, et al., 2014 Efficient Marker-Free Recovery of Custom Genetic Modifications with CRISPR/Cas9 in Caenorhabditis elegans. Genetics 198: 837–846. https://doi.org/10.1534/genetics.114.169730

2. Bone C. R., E. C. Tapley, M. Gorjanacz, and D. A. Starr, 2014 The Caenorhabditis elegans SUN protein UNC-84 interacts with lamin to transfer forces from the cytoplasm to the nucleoskeleton during nuclear migration. Mol Biol Cell 25: 2853–2865. https://doi.org/10.1091/mbc.e14-05-0971

3. Bone C. R., and D. A. Starr, 2016 Nuclear migration events throughout development. J Cell Sci 129: 1951–1961. https://doi.org/10.1242/jcs.179788

4. Bone C. R., Y.-T. Chang, N. E. Cain, S. P. Murphy, and D. A. Starr, 2016 Nuclei migrate through constricted spaces using microtubule motors and actin networks in C. elegans hypodermal cells. Development 143: 4193–4202. https://doi.org/10.1242/dev.141192

5. Brenner S., 1974 The genetics of Caenorhabditis elegans. Genetics.

6. Cabantous S., T. C. Terwilliger, and G. S. Waldo, 2005 Protein tagging and detection with engineered self-assembling fragments of green fluorescent protein. Nat Biotechnol 23: 102–107. https://doi.org/10.1038/nbt1044

7. Cain N. E., E. C. Tapley, K. L. McDonald, B. M. Cain, and D. A. Starr, 2014 The SUN protein UNC-84 is required only in force-bearing cells to maintain nuclear envelope architecture. J Cell Biology 206: 163–172. https://doi.org/10.1083/jcb.201405081

8. Cain N. E., Z. Jahed, A. Schoenhofen, V. A. Valdez, B. Elkin, et al., 2018 Conserved SUN-KASH Interfaces Mediate LINC Complex-Dependent Nuclear Movement and Positioning. Curr Biol 28: 3086–3097.e4. https://doi.org/10.1016/j.cub.2018.08.001

9. Chan E., and J. Nance, 2013 Mechanisms of CDC-42 activation during contact-induced cell polarization. J Cell Sci 126: 1692–1702. https://doi.org/10.1242/jcs.124594

10. Chang Y.-T., D. Dranow, J. Kuhn, M. Meyerzon, M. Ngo, et al., 2013 toca-1 Is in a Novel Pathway That Functions in Parallel with a SUN-KASH Nuclear Envelope Bridge to Move Nuclei in Caenorhabditis elegans. Genetics 193: 187–200. https://doi.org/10.1534/genetics.112.146589

11. Consortium C. elegans D. M., 2012 large-scale screening for targeted knockouts in the Caenorhabditis elegans genome. G3 Genes Genomes Genetics 2: 1415–1425. https://doi.org/10.1534/g3.112.003830

12. Cox E. A., and J. Hardin, 2004 Sticky worms: adhesion complexes in C. elegans. J Cell Sci 117: 1885–1897. https://doi.org/10.1242/jcs.01176

13. Davidson P. M., C. Denais, M. C. Bakshi, and J. Lammerding, 2014 Nuclear Deformability Constitutes a Rate-Limiting Step During Cell Migration in 3-D Environments. Cell Mol Bioeng 7: 293–306. https://doi.org/10.1007/s12195-014-0342-y

14. DeMaso C. R., I. Kovacevic, A. Uzun, and E. J. Cram, 2011 Structural and functional evaluation of C. elegans filamins FLN-1 and FLN-2. Plos One 6: e22428. https://doi.org/10.1371/journal.pone.0022428

15. Denais C. M., R. M. Gilbert, P. Isermann, A. L. McGregor, M. te Lindert, et al., 2016 Nuclear envelope rupture and repair during cancer cell migration. Science 352: 353–358. https://doi.org/10.1126/science.aad7297

16. Doonan R., J. Hatzold, S. Raut, B. Conradt, and A. Alfonso, 2008 HLH-3 is a C. elegans Achaete/Scute protein required for differentiation of the hermaphrodite-specific motor neurons. Mech Develop 125: 883–893. https://doi.org/10.1016/j.mod.2008.06.002

17. Evans T. C., 2006 Transformation and microinjection (April 6, 2006), WormBook, ed. The C. elegans Research Community, WormBook, doi/10.1895/wormbook.1.108.1.

18. Francis R., and R. H. Waterston, 1991 Muscle cell attachment in Caenorhabditis elegans. The Journal of cell biology 114: 465–479.

19. Fraser A. G., R. S. Kamath, P. Zipperlen, M. Martinez-Campos, M. Sohrmann, et al., 2000 Functional genomic analysis of C. elegans chromosome I by systematic RNA interference. Nature 408: 325–330. https://doi.org/10.1038/35042517

20. Fridolfsson H. N., N. Ly, M. Meyerzon, and D. A. Starr, 2010 UNC-83 coordinates kinesin-1 and dynein activities at the nuclear envelope during nuclear migration. Dev Biol 338: 237–250. https://doi.org/10.1016/j.ydbio.2009.12.004

21. Fridolfsson H. N., and D. A. Starr, 2010 Kinesin-1 and dynein at the nuclear envelope mediate the bidirectional migrations of nuclei. J Cell Biology 191: 115–128. https://doi.org/10.1083/jcb.201004118

22. Fridolfsson H. N., L. A. Herrera, J. N. Brandt, N. E. Cain, G. J. Hermann, et al., 2018 Genetic Analysis of Nuclear Migration and Anchorage to Study LINC Complexes During Development of Caenorhabditis elegans, p. 163–180 in The LINC Complex.

23. Frøkjær-Jensen C., M. W. Davis, C. E. Hopkins, B. J. Newman, J. M. Thummel, et al., 2008 Single-copy insertion of transgenes in Caenorhabditis elegans. Nat Genet 40: 1375–1383. https://doi.org/10.1038/ng.248

24. Frøkjær-Jensen C., M. W. Davis, M. Ailion, and E. M. Jorgensen, 2012 Improved Mos1-mediated transgenesis in C. elegans. Nat Methods 9: 117–118. https://doi.org/10.1038/nmeth.1865

25. Frøkjær-Jensen C., M. W. Davis, M. Sarov, J. Taylor, S. Flibotte, et al., 2014 Random and targeted transgene insertion in Caenorhabditis elegans using a modified Mos1 transposon. Nat Methods 11: 529–534. https://doi.org/10.1038/nmeth.2889

26. Fu Y., L. K. Chin, T. Bourouina, A. Q. Liu, and A. M. J. Vandongen, 2012 Nuclear deformation during breast cancer cell transmigration. Lab Chip 12: 3774–3778. https://doi.org/10.1039/c2lc40477j

27. Gensbittel V., M. Kräter, S. Harlepp, I. Busnelli, J. Guck, et al., 2021 Mechanical Adaptability of Tumor Cells in Metastasis. Dev. Cell 56: 164–179. https://doi.org/10.1016/j.devcel.2020.10.011

28. Giuliani C., F. Troglio, Z. Bai, F. B. Patel, A. Zucconi, et al., 2009 Requirements for F-BAR proteins TOCA-1 and TOCA-2 in actin dynamics and membrane trafficking during Caenorhabditis elegans oocyte growth and embryonic epidermal morphogenesis. Plos Genet 5: e1000675. https://doi.org/10.1371/journal.pgen.1000675

29. Hall D. H., and Z. F. Altun, 2008 C. Elegans Atlas. Cold Spring Harbor Laboratory Pr.

30. Hao H., S. Kalra, L. E. Jameson, L. A. Guerrero, N. E. Cain, et al., 2021 The Nesprin-1/-2 ortholog ANC-1 regulates organelle positioning in C. elegans independently from its KASH or actin-binding domains. Elife 10: e61069. https://doi.org/10.7554/elife.61069

31. Ho H.-Y. H., R. Rohatgi, A. M. Lebensohn, M. Le, J. Li, et al., 2004 Toca-1 mediates Cdc42-dependent actin nucleation by activating the N-WASP-WIP complex. Cell 118: 203–216. https://doi.org/10.1016/j.cell.2004.06.027

32. Ho J., V. A. Valdez, L. Ma, and D. A. Starr, 2018 Characterizing Dynein’s Role in P-cell Nuclear Migration using an Auxin-Induced Degradation System. MicroPublication:Biology. https://doi.org/10.17912/w2w96j

33. Ho J., L. A. Guerrero, D. Libuda, G. G. Luxton, and D. A. Starr, 2023 A CDC-42-regulated actin network is necessary for nuclear migration through constricted spaces in C. elegans. bioRxiv 2023.06.22.546138. https://doi.org/10.1101/2023.06.22.546138

34. Huang X., H.-J. Cheng, M. Tessier-Lavigne, and Y. Jin, 2002 MAX-1, a Novel PH/MyTH4/FERM Domain Cytoplasmic Protein Implicated in Netrin-Mediated Axon Repulsion. Neuron 34: 563–576. https://doi.org/10.1016/s0896-6273(02)00672-4

35. Jahed Z., H. Hao, V. Thakkar, U. T. Vu, V. A. Valdez, et al., 2019 Role of KASH domain lengths in the regulation of LINC complexes. Mol Biol Cell mbcE19020079. https://doi.org/10.1091/mbc.e19-02-0079

36. Junt T., H. Schulze, Z. Chen, S. Massberg, T. Goerge, et al., 2007 Dynamic visualization of thrombopoiesis within bone marrow. Science 317: 1767–1770. https://doi.org/10.1126/science.1146304

37. Kamath R., A. Fraser, Y. Dong, G. Poulin, R. Durbin, et al., 2003 Systematic functional analysis of the Caenorhabditis elegans genome using RNAi. Nature 421: 231–237. https://doi.org/10.1038/nature01278

38. Kim H.-S., R. Murakami, S. Quintin, M. Mori, K. Ohkura, et al., 2011 VAB-10 spectraplakin acts in cell and nuclear migration in Caenorhabditis elegans. Development 138: 4013–4023. https://doi.org/10.1242/dev.059568

39. Kovacevic I., J. M. Orozco, and E. J. Cram, 2013 Filamin and Phospholipase C-ε Are Required for Calcium Signaling in the Caenorhabditis elegans Spermatheca, (A. D. Chisholm, Ed.). Plos Genet 9: e1003510. https://doi.org/10.1371/journal.pgen.1003510

40. Malone C. J., W. D. Fixsen, H. R. Horvitz, and M. Han, 1999 UNC-84 localizes to the nuclear envelope and is required for nuclear migration and anchoring during C. elegans development. Development 126: 3171–3181. https://doi.org/10.1242/dev.126.14.3171

41. McGee M. D., R. Rillo, A. S. Anderson, and D. A. Starr, 2006 UNC-83 IS a KASH protein required for nuclear migration and is recruited to the outer nuclear membrane by a physical interaction with the SUN protein UNC-84. Mol Biol Cell 17: 1790–1801. https://doi.org/10.1091/mbc.e05-09-0894

42. McGregor A. L., C.-R. Hsia, and J. Lammerding, 2016 Squish and squeeze—the nucleus as a physical barrier during migration in confined environments. Curr Opin Cell Biol 40: 32–40. https://doi.org/10.1016/j.ceb.2016.01.011

43. McIntire S. L., R. J. Reimer, K. Schuske, R. H. Edwards, and E. M. Jorgensen, 1997 Identification and characterization of the vesicular GABA transporter. Nature 389: 870–876. https://doi.org/10.1038/39908

44. Meyerzon M., H. N. Fridolfsson, N. Ly, F. J. McNally, and D. A. Starr, 2009 UNC-83 is a nuclear-specific cargo adaptor for kinesin-1-mediated nuclear migration. Development 136: 2725-2733. https://doi.org/10.1242/dev.038596

45. Mirdita M., K. Schütze, Y. Moriwaki, L. Heo, S. Ovchinnikov, et al., 2022 ColabFold: making protein folding accessible to all. Nat. Methods 19: 679–682. https://doi.org/10.1038/s41592-022-01488-1

46. Nakamura F., T. P. Stossel, and J. H. Hartwig, 2011 The filamins: organizers of cell structure and function. Cell Adhes Migr 5: 160–169. https://doi.org/10.4161/cam.5.2.14401

47. Olmos Y., 2022 The ESCRT Machinery: Remodeling, Repairing, and Sealing Membranes. Membranes 12: 633. https://doi.org/10.3390/membranes12060633

48. Paix A., A. Folkmann, and G. Seydoux, 2017 Precision genome editing using CRISPR-Cas9 and linear repair templates in C. elegans. Methods 121: 86–93. https://doi.org/10.1016/j.ymeth.2017.03.023

49. Pettersen E. F., T. D. Goddard, C. C. Huang, E. C. Meng, G. S. Couch, et al., 2021 UCSF ChimeraX: Structure visualization for researchers, educators, and developers. Protein Sci. 30: 70–82. https://doi.org/10.1002/pro.3943

50. Raab M., M. Gentili, H. de Belly, H. R. Thiam, P. Vargas, et al., 2016 ESCRT III repairs nuclear envelope ruptures during cell migration to limit DNA damage and cell death. Science 352: aad7611 362. https://doi.org/10.1126/science.aad7611

51. Razinia Z., T. Mäkelä, J. Ylänne, and D. A. Calderwood, 2012 Filamins in Mechanosensing and Signaling. Annu Rev Biophys 41: 227–246. https://doi.org/10.1146/annurev-biophys-050511-102252

52. Renkawitz J., A. Kopf, J. Stopp, I. de Vries, M. K. Driscoll, et al., 2019 Nuclear positioning facilitates amoeboid migration along the path of least resistance. Nature 42: 7. https://doi.org/10.1038/s41586-019-1087-5

53. Riedl J., A. H. Crevenna, K. Kessenbrock, J. H. Yu, D. Neukirchen, et al., 2008 Lifeact: a versatile marker to visualize F-actin. Nat Methods 5: 605–607. https://doi.org/10.1038/nmeth.1220

54. Salvermoser M., D. Begandt, R. Alon, and B. Walzog, 2018 Nuclear Deformation During Neutrophil Migration at Sites of Inflammation. Front. Immunol. 9: 2680. https://doi.org/10.3389/fimmu.2018.02680

55. Sarin S., V. Bertrand, H. Bigelow, A. Boyanov, M. Doitsidou, et al., 2010 Analysis of multiple ethyl methanesulfonate-mutagenized Caenorhabditis elegans strains by whole-genome sequencing. Genetics 185: 417-430. https://doi.org/10.1534/genetics.110.116319

56. Seyfried T. N., and L. C. Huysentruyt, 2012 On the Origin of Cancer Metastasis. Crit. Rev. Oncog. 18: 43–73. https://doi.org/10.1615/critrevoncog.v18.i1-2.40

57. Shemiakina I. I., G. V. Ermakova, P. J. Cranfill, M. A. Baird, R. A. Evans, et al., 2012 A monomeric red fluorescent protein with low cytotoxicity. Nat Commun 3: 1204. https://doi.org/10.1038/ncomms2208

58. Shi L., Y. Jian, M. Li, T. Hao, C. Yang, et al., 2022 Filamin FLN-2 promotes MVB biogenesis by mediating vesicle docking on the actin cytoskeleton. J Cell Biol 221: e202201020. https://doi.org/10.1083/jcb.202201020

59. Shin J.-W., K. R. Spinler, J. Swift, J. A. Chasis, N. Mohandas, et al., 2013 Lamins regulate cell trafficking and lineage maturation of adult human hematopoietic cells. Proc National Acad Sci 110: 18892–18897. https://doi.org/10.1073/pnas.1304996110

60. Starr D. A., G. J. Hermann, C. J. Malone, W. Fixsen, J. R. Priess, et al., 2001 unc-83 encodes a novel component of the nuclear envelope and is essential for proper nuclear migration. Development 128: 5039–5050. https://doi.org/10.1242/dev.128.24.5039

61. Starr D. A., and H. N. Fridolfsson, 2010 Interactions between nuclei and the cytoskeleton are mediated by SUN-KASH nuclear-envelope bridges. Annu Rev Cell Dev Bi 26: 421–444. https://doi.org/10.1146/annurev-cellbio-100109-104037

62. Starr D. A., 2019 A network of nuclear envelope proteins and cytoskeletal force generators mediates movements of and within nuclei throughout Caenorhabditis elegans development. Exp Biol Med 244: 1323–1332. https://doi.org/10.1177/1535370219871965

63. Stiernagle T., 2006 Maintenance of C. elegans (February 11, 2006), WormBook, ed. The C. elegans Research Community, WormBook, doi/10.1895/wormbook.1.101.1.

64. Sulston J. E., and H. R. Horvitz, 1977 Post-embryonic cell lineages of the nematode, Caenorhabditis elegans. Dev Biol 56: 110–156. https://doi.org/10.1016/0012-1606(77)90158-0

65. Sulston J. E., and H. R. Horvitz, 1981 Abnormal cell lineages in mutants of the nematode Caenorhabditis elegans. Dev Biol 82: 41–55. https://doi.org/10.1016/0012-1606(81)90427-9

66. Thiam H.-R., P. Vargas, N. Carpi, C. L. Crespo, M. Raab, et al., 2016 Perinuclear Arp2/3-driven actin polymerization enables nuclear deformation to facilitate cell migration through complex environments. Nat Commun 7: 10997. https://doi.org/10.1038/ncomms10997

67. Williams-Masson E. M., A. N. Malik, and J. Hardin, 1997 An actin-mediated two-step mechanism is required for ventral enclosure of the C. elegans hypodermis. Development (Cambridge, England) 124: 2889–2901.

68. Xia Y., I. L. Ivanovska, K. Zhu, L. Smith, J. Irianto, et al., 2018 Nuclear rupture at sites of high curvature compromises retention of DNA repair factors. J. Cell Biol. 217: 3796–3808. https://doi.org/10.1083/jcb.201711161

